# Predicting mutation-rate variation across the genome using epigenetic data

**DOI:** 10.64898/2026.01.30.702885

**Authors:** Machiko Katori, Tetsuya J. Kobayashi, Magnus Nordborg, Shoi Shi

## Abstract

Mutation rate variation is a fundamental driver of evolution, yet how it is locally patterned across genomes and structured by chromatin context remains unresolved. Here, we integrate genome-wide profiles of histone marks, DNA methylation and chromatin accessibility in *Arabidopsis thaliana* with de novo mutation data to model mutation probability at the level of coding sequence (CDS). Using non-negative matrix factorization, we identify 15 combinatorial epigenetic patterns whose graded mixtures stratify CDSs into six classes with distinct mutation probabilities. A generalized linear model based on pattern weights predicts local mutation probability and outperforms models based on sequence context, expression and classical genomic categories. These patterns capture context-dependent variation that is obscured by gene-level summaries and single-feature analyses. Cluster-level differences are partly retained in mutation-accumulation lines, indicating persistence into heritable mutational input. Under hypoxia, stress-responsive chromatin remodeling redistributes epigenetic contexts associated with higher predicted mutation probability toward hypoxia-responsive genes and DNA-repair pathways. Together, our results provide a CDS-resolved and interpretable framework linking combinatorial epigenomic context to mutational input, clarifying how dynamic chromatin states shape local mutation-rate heterogeneity.

## Introduction

Mutations provide the raw material for evolution, yet are mostly deleterious and sophisticated mechanisms have evolved to avoid mutation [1]. That mutation rates vary greatly between species is well established, but they are now also recognized to be highly nonuniform across genomes, varying with local context shaped by both static sequence features and dynamic properties of genome organization that can shift with environmental or developmental cues (e.g., replication timing and chromatin organization) [2, 3, 4].

This raises a key question: does mutation-rate heterogeneity across genomes contradict the classical evolutionary view [5] that mutation is random with respect to fitness? Genome-wide studies across taxa show that epigenomic states can modulate the efficiency of DNA repair. In human cells, H3K36me3 promotes recruitment of the mismatch repair factor MSH6 [6]. In *Arabidopsis thaliana*, H3K4me1 has likewise been linked to preferential recruitment of mismatch repair [7]. Consistent with these repair-associated marks, regions enriched for H3K36me3 (exons and actively expressed genes) [8, 9] and for H3K4me1 (essential genes) [10] are associated with lower de novo mutation rates. However, category-level differences in mutation rates — typically inferred from comparisons between broad genomic or functional categories — can arise from context-dependent processes of DNA damage and repair. Thus, they need not indicate adaptive targeting of mutational input [11]. Moreover, such comparisons are sensitive to technical biases (e.g., mappability, repetitiveness, coverage, and category-dependent error profiles of variant callings) [12, 13, 14] and to differences in selective constraint [15]. Together, these considerations motivate analyses that resolve mutation-rate heterogeneity at finer scales, as a prerequisite for interpreting whether epigenome-associated patterns reflect nonuniformity alone or also have fitness-relevant implications.

One straightforward strategy is to restrict attention to a single genomic class, coding sequences (CDSs). This within-class design allows us to investigate heterogeneity in mutation rates with fewer confounders while retaining interpretability given well-defined CDS annotations. Yet CDS-level epigenetic patterns can be partially obscured by chromatin organization [3, 4, 16] and broad chromatin-state annotations (e.g., ChromHMM) [17, 18, 19]. What remains lacking is a CDS-resolved map of epigenetic configurations that can account for local mutation-rate heterogeneity beyond these coarse stratifications.

Here, using extensive epigenomic datasets in *A. thaliana* together with de novo mutation data, we systematically analyze local epigenetic patterns within CDSs and quantify their contribution to the observed heterogeneity in mutation rates. We identify CDS-resolved epigenetic configurations associated with distinct local mutation-rate regimes, providing a high-resolution view of the local mutational landscape. Informed by environmentally responsive epigenomic data, we further raise the possibility that regulatory plasticity under environmental stress can shift epigenetic contexts toward configurations with higher predicted mutation probability at fitness-relevant CDSs, with potential consequences for long-term genetic adaptation.

## Results

### Combinatorial epigenetic patterns at coding-sequence resolution

Assessing the relationship between epigenomic features and mutation probability is hindered by category-specific biases in variant calling [12, 13, 14] and epigenetic profiling [20, 21]. In addition, single-mark associations are limited because DNA damage and repair integrate combinations of cues, and many epigenomic tracks are collinear. We therefore adopted an analysis framework to address both issues. We restricted analyses to coding sequences (CDSs), where single-nucleotide variants can be called with high confidence [12] and epigenomic profiles are comparatively stable. We then learned non-negative, additive modules of co-occurring features from a matrix of histone marks [22], DNA methylation [23], chromatin accessibility [10], and GC content. non-negative matrix factorization (NMF) yields graded mixture weights per segment (Fig. 1a; Supplementary Fig. 1a–d), avoids sign cancellation and stabilizes inference relative to raw features (Supplementary Fig. 1e, f). The resulting combinatorial 15 patterns (Fig. 1b; Supplementary Fig. 1g–j) recapitulate known repressive constellations [24] and reveal GC-coupled states not captured by classical annotations, providing a low-dimensional epigenetic basis at CDS resolution. We next asked whether differences in these epigenetic mixtures stratify mutation probability.

**Fig. 1.**
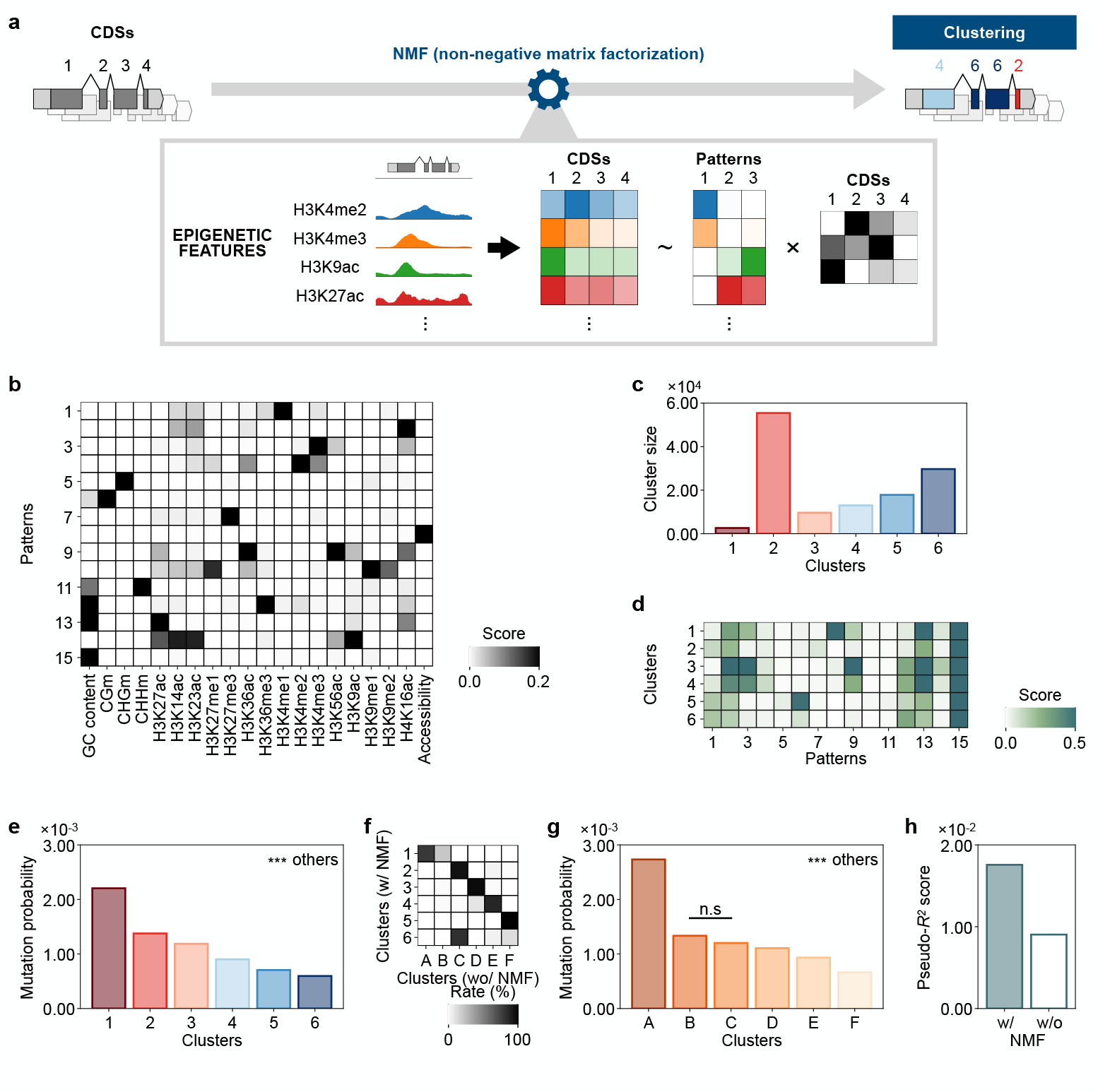
Combinatorial epigenetic patterns define six CDS clusters with distinct mutation probabilities. **(a)** Overview of the workflow for epigenetic analysis. **(b)** Epigenetic pattern matrix extracted by NMF. **(c)** Cluster size. **(d)** Heatmap showing the average epigenetic pattern weights across clusters. **(e)** Mutation probabilities of clusters. For all cluster pairs, the log-likelihood ratio test yielded \*\*\**p <* 0.001 after Bonferroni correction. **(f)** Heatmap showing the transition from the original clusters obtained with on NMF (clusters 1-6) to those obtained without NMF (clusters A-F). **(g)** Mutation probabilities of clusters defined without NMF. For all cluster pairs, unless otherwise noted (n.s., not significant), the log-likelihood ratio test yielded \*\*\**p <* 0.001 after Bonferroni correction. **(h)** Pseudo-*R*^2^ score for mutation probabilities explained by the six clusters defined with or without NMF.

### Six CDS clusters show distinct mutation probabilities

Pattern mixtures revealed six clusters of CDSs with distinct epigenomic combinations (Fig. 1c, d; Supplementary Fig. 2a–c). CDSs from different clusters often co-occurred within the same gene, exposing sub-gene organization (Supplementary Fig. 2d). Mutation probabilities estimated from de novo somatic mutations [10] differed statistically among clusters (Fig. 1e). By contrast, clustering based on raw epigenomic features weakened these differences and eliminated one cluster-level contrast (Fig. 1f, g), reducing the explanatory power of cluster assignments for mutation-rate variability (Fig. 1h). Stable cluster stratification required fitting NMF with a sufficiently large number of patterns; however the clustering assignments were robust once the representation was adequately learned (Supplementary Fig. 3). ChromHMM-based states [20, 21] showed limited overlap with our CDS clusters (Supplementary Fig. 4). This is consistent with ChromHMM learning genome-wide chromatin-state segmentation [17, 19], thus capturing a signal distinct from the CDS-specific combinatorial signatures identified here.

### Classical genomic categories do not explain these differences

We asked whether classical genomic and functional categories could account for the cluster-level separation in mutation probability. CDSs were stratified by length, distance to the transcription start site (TSS), chromosomal position, expression level, essentiality and lethality (Fig. 2a; Supplementary Fig. 5a). Cluster-level differences persisted across strata; modest trends associated with CDS length and TSS distance were complementary rather than redundant with cluster identity (Fig. 2b, c; Supplementary Fig. 5b). Stratifying CDSs by expression level reduced the explanatory power of cluster identity for mutation probability. However, the relative ordering of clusters was preserved within expression strata (Fig. 2d), indicating that expression alone does not account for the observed separation, despite potential links between transcription-associated processes, DNA repair and epigenetic features [25, 26]. Consistent with this, expression levels showed similar distributions across the six clusters (Supplementary Fig. 5c), supporting the added value of CDS-resolved epigenomic stratification. Dividing CDSs based on essentiality or lethality also reduced overall explanatory power, yet the cluster-level differences remained evident even within non-essential or non-lethal subsets (Supplementary Fig. 5d–f). Synonymous fractions were similar across clusters (Supplementary Fig. 5g, h), and genes dominated by a single cluster showed no consistent gene ontology enrichment (Supplementary Fig. 6). Together, these observations indicate that the observed separation reflects epigenetic patterning rather than classical functional categories or obvious proxies of selective constraint.

**Fig. 2.**
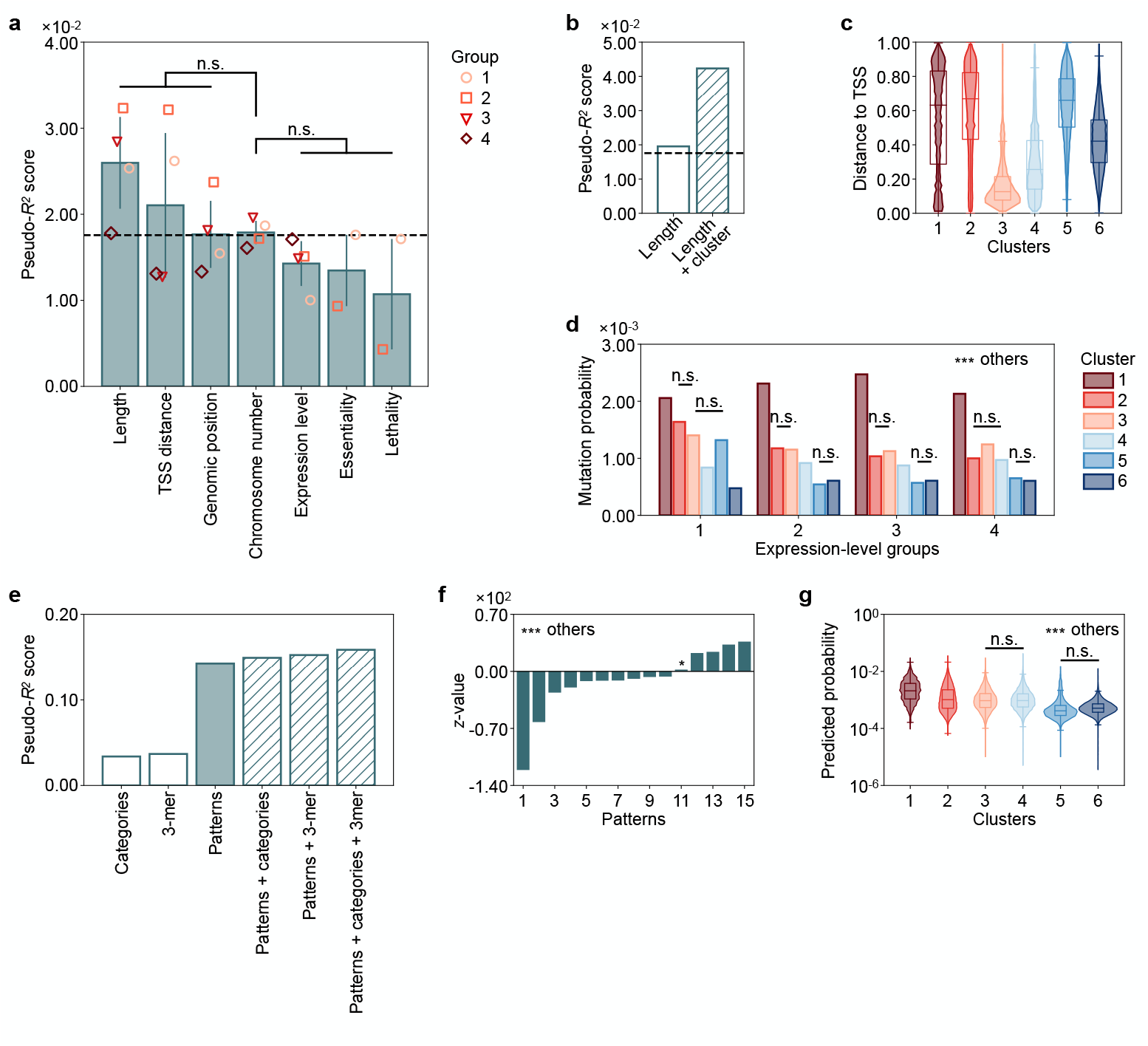
Pattern weights predict local mutation probabilities over classical genomic categories. **(a)** Pseudo-*R*^2^ score for mutation probabilities explained by the six clusters within each group stratified by classical genomic categories. Student’s *t*-test after Bonferroni correction on the basal chromosomal number stratification shows no statistical significance compared with the other categories. **(b)** Pseudo-*R*^2^ score for mutation probabilities explained by four length-based groups and by the combination of length-based groups and the six clusters. **(c)** Distribution of exon positions across clusters. **(d)** Mutation probabilities of clusters calculated for each expression group. For all cluster pairs within each expression group, unless otherwise noted (n.s., not significant), the log-likelihood ratio test yielded \*\*\**p <* 0.001 after Bonferroni correction. **(e)** Pseudo-*R*^2^ score for mutation probabilities explained by various combinations of classical genomic categories, 3-mer, and epigenetic patterns. **(f)** *z*-values of the epigenetic patterns in the GLM prediction. For all patterns, unless otherwise noted (\**p <* 0.05), the Wald test yielded \*\*\**p <* 0.001. **(g)** Distribution of predicted probabilities among clusters. For all pairs of Clusters *i* and *j* (*i < j*), unless otherwise noted (n.s., not significant), a one-sided Mann-Whitney *U* test yielded \*\*\**p <* 0.001 after Bonferroni correction.

### Pattern weights predict local mutation probability

To quantify effects at segment resolution, we fit a generalized linear model with pattern weights as predictors. Despite substantial sparsity in the response (91.5% of CDSs lacked observed mutations), the model captured a robust signal and outperformed models based on classical categories or sequence context; adding those covariates provided no further gain (Fig. 2e; Supplementary Fig. 7a–c). All 15 patterns contributed to the model (Fig. 2f). A pattern enriched for H3K4me1 (pattern 1) showed the strongest negative association, consistent with previous report linking H3K4me1 to reduced mutation via mismatch-repair targeting in plants [7]. Patterns that shared features sometimes had opposite effects, indicating context-dependent interactions rather than simple additivity. Predicted mutation probabilities recapitulated the empirical differences among clusters (Fig. 2g; Supplementary Fig. 7d).

### Epigenome-linked variation in mutation probability at fitness-relevant loci

We asked whether epigenome-linked differences in mutation probability are detectable in heritable mutations and at fitness-relevant loci. We used mutation-accumulation (MA) lines [15] to test persistence across generations and hypoxia-treated plants [27] to examine stress-driven reweighting of epigenomic contexts. In MA lines, cluster-level separation was partly retained: CDSs in clusters 5 and 6 showed statistically lower mutation probabilities than those in clusters 1 and 2 (Fig. 3a; Supplementary Fig. 8a). These observations suggest that epigenome-linked differences identified from somatic mutations can persist across generations, supporting their relevance for evaluating potential fitness consequences.

**Fig. 3.**
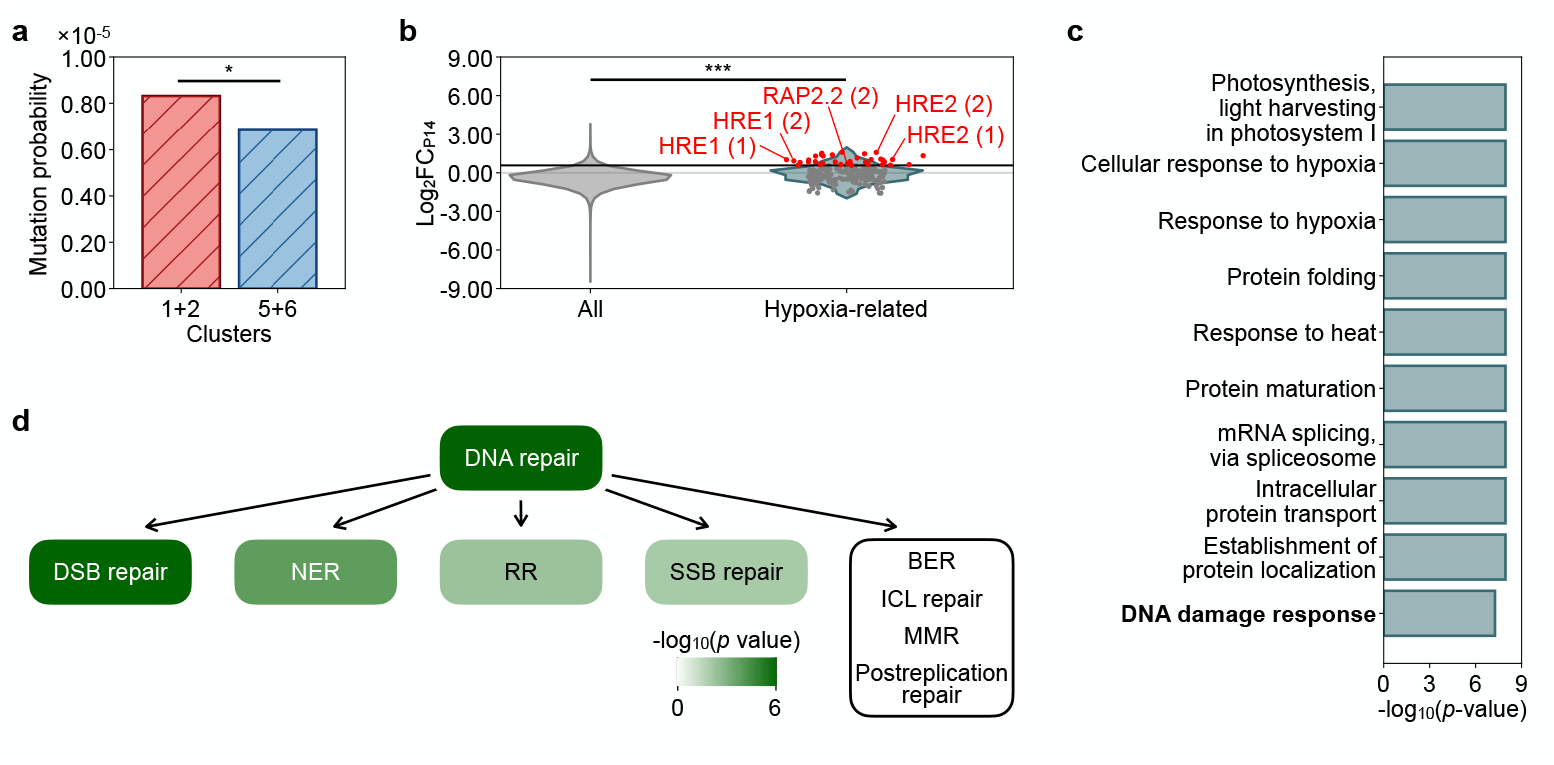
Epigenome-linked variation in mutation probability at fitness-relevant loci. **(a)** MA line-based mutation probabilities of CDSs within clusters 1 and 2 and within clusters 5 and 6. The log-likelihood ratio test yielded \*\*\**p <* 0.05 after Bonferroni correction. **(b)** Distribution of pattern-14 activation scores for all CDSs and for CDSs from hypoxia-related genes, with data points in the top 5% highlighted in red. Selected points are labeled with gene names, and the numbers indicate the CDS order from TSS. A one-sided Mann-Whiney *U* test revealed *∗∗∗p <* 0.001. **(c)** Top 10 GO terms identified by GSEA, ranked by the maximum pattern-14 activation score per gene. **(d)** Enrichment analysis based on the Mann-Whitney *U* test *p*-values after Bonferroni correction for CDSs from genes belonging to each GO term compared with the background distribution of pattern-14 activation. Green shading indicates *p*-values, while GO terms in the white area show no significant enrichment (*p >* 0.05).

We then asked whether environmental stress redistributes epigenomic signatures associated with higher predicted mutation probability across CDSs. Under hypoxia, a simple pattern-14 activation score (capturing co-induction of H3K9ac and H3K14ac and associated with higher predicted mutation probability; Supplementary Information) increased at hypoxia-responsive genes relative to the genomic background (Fig. 3b; Supplementary Fig. 8b, c). Among the 53 hypoxia-responsive genes [27, 28], 22 contained at least one CDS within the genome-wide top 5% of activation scores. This activation score was only weakly correlated with transcriptional induction (Pearson’s *r* = 0.20), indicating that it captures information complementary to expression. Notably, this set included hypoxia-tolerance loci with gain-of-function alleles, namely HRE1, HRE2 and PAR2.2 [29, 30], which occupied the extreme tail of the activation-score distribution even when their transcriptional induction was not comparably extreme. Gene-level enrichment analysis recovered hypoxia-related terms for both activation-score and expression-based rankings, but DNA-repair pathways were more strongly enriched in activation-score ranking (Fig. 3c; Supplementary Fig. 8d). This enrichment was consistent at CDS resolution with multiple DNA-repair subpathways showing significant enrichment (Fig. 3d; Supplementary Fig. 8e). Together, these results support a model in which stress-driven epigenomic reconfiguration is associated with elevated predicted mutation probability at fitness-relevant CDSs under stress, including hypoxia-tolerance genes and DNA-repair pathways, with potentially positive consequences for adaptation.

## Discussion

Heterogeneity of mutation rates across the genome reflects processes operating at multiple scales, from higher-order chromatin organization to local determinants of DNA damage and repair [2, 3, 4, 6, 7, 31, 32, 33]. Yet evolutionary interpretation remains complicated by mismatched resolution across studies [34] and is further confounded by technical artifacts and natural selection, particularly when mutation rates are compared across broad functional categories [12, 13, 14]. Here, we developed an analysis that models combinations of epigenomic features at the level of individual coding sequences (CDSs) using non-negative matrix factorization (Fig. 1a). This pattern-based approach resolved 15 epigenetic patterns (Fig. 1b) and partitioned CDSs into six clusters with distinct mutation probabilities (Fig. 1c–e). This combinatorial view accounts for more of the variation in local mutation probability than single-feature analyses (Fig. 1f–h), classical genomic categories, or sequence context (Fig. 2), revealing an additional axis of mutational heterogeneity within functionally matched regions. Together, these results provide a CDS-resolved view of mutational heterogeneity at a finer resolution than previously appreciated.

Our CDS-resolved, pattern-based view suggests mechanistic hypotheses for how chromatin configurations modulate mutation probability (Fig. 1b; Fig. 2f). The negative association of H3K4me1-rich pattern with mutation probability aligns with preferential engagement of mismatch repair on H3K4-decorated chromatin [7]. More broadly, our results expose mutation-probability structure that is obscured by gene-level summaries and single-feature analyses. A particularly informative case concerns transcription: evidence for a simple transcription-mutation relationship remains mixed [35], likely reflecting opposing effects of transcription-coupled activity on repair (e.g., transcription-coupled repair) and vulnerability [26, 36]. In our data, acetylation-rich contexts, which are often associated with open and transcription-permissive chromatin, showed opposing associations: H3K14ac combined with H4 acetylation predicted lower mutation probability, whereas co-occurring H3K9ac-H3K14ac predicted higher probability. These opposing associations are compatible with a model in which the balance between repair and vulnerability depends on the specific co-mark configuration. Supporting this interpretation, the six CDS clusters exhibited similar expression levels (Supplementary Fig. 5c), and their differences in mutation probability were not fully explained by expression level (Fig. 2d). Notably, this suggests that mutation-probability heterogeneity within CDSs can be stratified beyond expression-linked chromatin states defined by ChromHMM [17, 18, 19]. Likewise, GC content showed context-dependent effects when paired with other marks, reinforcing that single-mark heuristics are insufficient and that combinatorial grammars better capture how chromatin state routes damage and repair. These hypotheses can be tested with epigenome editing [37] that recruits individual marks versus specific co-mark pairs at endogenous CDSs, coupled to longitudinal readouts of de novo mutations and targeted perturbation of repair factors. Taken together, our CDS-resolved framework provides a higher-resolution basis for evaluating whether mutational input is independent of fitness, and for assessing the potential contribution of epigenomic context.

A central question is whether context-dependent mutational heterogeneity has consequences for fitness. In mutation-accumulation lines [15], the cluster-level differences we observed in somatic mutations were partly retained across generations (Fig. 3a), suggesting that cluster-associated biases are not purely somatic but reflect germ-line mutation rates as well. Synonymous variant fractions (Supplementary Fig. 5g) and gene-level enrichment (Supplementary Fig. 6) were comparable across clusters, arguing against obvious differences in selective constraint. This concordance contrasts with human studies in which germline mutational heterogeneity appears to represent a subset of somatic heterogeneity [38, 39]. One plausible explanation is that, unlike mammals, *A. thaliana* lacks an early-segregated germline, blurring the distinction between somatic and heritable mutational processes [40, 41]. Stress remodeling provides an additional lens. Under hypoxia, joint induction of H3K9ac and H3K14ac (pattern-14 activation) was elevated in CDSs of hypoxia-responsive genes (Fig. 3b) and was only weakly correlated with transcriptional up-regulation (Supplementary Fig. 8c, d). Crucially, loci with known gain-of-function alleles conferring hypoxia tolerance, such as HRE1, HRE2, and PAR2.2 [29, 30], fell among CDSs with the highest pattern-14 activation scores (Fig. 3b). Together, these observations raise the possibility that stress-driven epigenomic reconfiguration could specifically increase mutation rates in genes with immediate relevance for adaptation. Whether such response would be adaptive has been hotly debated [11, 42]. Interestingly, pattern-14 activation was also enriched in DNA repair pathways (Fig. 3c, d), suggesting that stress may increase mutational input not only at putatively adaptive targets but also within genomestability modules. Such shifts would of course further expand the overall mutational input available for selection.

However, if stress-associated redistribution of mutational input has fitness-relevant consequences, a key question is whether a mechanism that modulates mutation probability in an epigenetic-context-dependent manner can be maintained by selection. A directly adaptive explanation is constrained by the drift-barrier hypothesis: selection on mutationrate modifiers is limited by the fitness consequences of the mutational-input changes they induce [43]. In our case, cluster-to-cluster differences in mutation probability are on the order of 10^*−*3^-10^*−*5^, depending on the dataset (Fig. 1e; Fig. 3a), making it nontrivial to explain how chromatin-dependent modulation of mutation probability could be maintained as an adaptive mechanism. An important consideration is that *A. thaliana* life history may alter these expectations: given the lack of early-segregated germline [40, 41], somatic lineages can in principle contribute to gamete formation, allowing selection to act on mutation-rate modifiers through both heritable mutations and somatic variation that contributes to reproduction. In addition, mutation-rate modulation may persist as a pleiotropic consequence of epigenome-based plasticity and stress-responsive chromatin remodeling, even if it is not itself a primary target of selection. Assessing whether these routes can plausibly account for the stress-associated redistribution of mutational input toward fitness-relevant loci will require explicit evolutionary models that incorporate plant life history, lack of early-segregated germline, the timing of mutational events, and the joint effects of stress responses on phenotype and genome stability [44].

More generally, it will be important to test how broadly such effects generalize across eukaryotes, given that core chromatin architectures and many DNA repair pathways are deeply conserved [45, 46, 47, 48]. This conservation is consistent with the possibility that combinatorial epigenomic conditioning of mutation probability represents a general principle, while the specific grammars of mark combinations and the stress programs that mobilize them are likely lineage-dependent. Comparative studies — for example, plants under abiotic stress, fungi under nutrient limitation, and mammalian cells under metabolic or inflammatory challenge — will help disentangle universal rules from lineagespecific strategies. In this broader view, epigenomic context emerges as a bridge between transient cellular states and long-term genetic outcomes, explaining how genomes both preserve stability and generate the variation on which evolution depends.

## Acknowledgements

We express our gratitude to the members of the Administrative Department at IIIS for their continuous support. We are grateful to Grey J. Monroe for valuable discussions related to epigenomic and mutation-rate analyses, and to Tetsuya Higashiyama, Takashi Tsuchimatsu, Haruka Ozaki, and Taro Toyoizumi for constructive comments and perspectives. We additionally thank Tetsuya Yamada for helpful input, and Yoko Toyoshima and Junko Kanoh for fruitful discussions during the ACT-X grant meeting. We acknowledge all members of the Shi laboratory, Kobayashi laboratory, and Nordborg laboratory for their contributions to discussions and the broader research environment.

This work was supported by the Japan Society for the Promotion of Science (JSPS) Grants-in-Aid for Scientific Research (KAKENHI) (22K21352 to MK and MN; 20H05894, 20H05903, 21K15136, 22K21351, 23H02518A, 23H02663, and 23K18147 to S.S.; 25H01365 to T.J.K.); the JSPS Research Fellowship (22KJ0931 and 25KJ0029 to M.K.); JST-CREST (JPMJCR24T4 and JPMJCR2551 to S.S.; PMJCR2011 and JPMJCR25Q2 to T.J.K.); JST ACT-X (JPMJAX2326 to M.K.); the World Premier International Research Center Initiative (WPI) from the Ministry of Education, Culture, Sports, Science and Technology (MEXT) to S.S. (WPI-IIIS); the Top Runners in Strategy of Transborder Advanced Researches (TRiSTAR) by MEXT to S.S.; the Japan Agency for Medical Research and Development (AMED) (JP21zf0127005 to S.S.), and the ANRI Scholarship to M.K.

## Author contributions

Study design, manuscript writing, and figure preparation were jointly carried out by MK and SS. MK conducted all data analyses under the guidance of SS. TJK and NM provided in-depth discussions and contributed significantly to the interpretation of the results. All authors carefully read the manuscript and provided comments.

