## Supplementary Information for "Predicting mutation-rate variation across the genome using epigenetic data"

**The PDF file includes:**

Methods
Supplementary Fig. 1 to Supplementary Fig. 8
Supplementary References

### Methods

#### Data acquisition

##### CDS regions

To minimize the effects of isoforms on our analysis, we focused on coding genes listed in the TAIR10 representative gene models. According to TAIR10 annotations, each gene is divided into the 5' untranslated region (5'UTR), coding sequence (CDS), intron, and 3'UTR. Throughout this study, we used CDSs as the primary unit of analysis. The relative position of each CDS was calculated as the distance from the midpoint of the CDS to the 5'UTR, normalized by the total gene length (from 5'UTR to 3'UTR).

##### Epigenetic features

DNA methylation data were obtained from GSM1085222, generated via Methyl-Seq as a part of the 10001 epigenome project [23]. Methylation levels were classified into CG, CHG, and CHH contexts (denoted CGm, CHGm, and CHHm, respectively) according to the TAIR10 reference genome. To construct a high-resolution landscape of histone modifications, we downloaded 64 ChIP-seq BigWig tracks from the Plant Chromatin State Database (PCSD) [22]. These tracks corresponded to the following histone marks: H3K27ac ($n = 2$), H3K14ac ($n = 1$), H3K23ac ($n = 5$), H3K27me1 ($n = 1$), H3K27me3 ($n = 7$), H3K36ac ($n = 2$), H3K36me3 ($n = 1$), H3K4me1 ($n = 6$), H3K4me2 ($n = 6$), H3K4me3 ($n = 13$), H3K56ac ($n = 1$), H3K9ac ($n = 5$), H3K9me1 ($n = 1$), H3K9me2 ($n = 12$), and H4K16ac ($n = 1$). When multiple replicates were available for a given mark, we used the average signal across replicates. In addition, chromatin accessibility data from ATAC-seq were obtained from published datasets [10].

##### Single-nucleotide variants

We utilized published somatic mutation data derived from mutation accumulation lines based on branch based on branch structures [50], which were reanalyzed to enhance the sensitivity of mutation detection [10]. In the original dataset, some insertion-deletion variants were mislabeled as single-nucleotide variants (SNVs); we corrected these annotations and used the curated SNV set for downstream analyses.

##### Expression data

Normalized RNA-seq expression data were obtained from GSE80744 and averaged across accessions [23]. Prior to analysis, expression values were transformed using $\log_{10}(1 + x)$. Genes lacking expression data were excluded from all downstream analyses, resulting in 128263 CDSs. The definitions of essential and lethal genes followed those described by Monroe et al. [10].

#### NMF pipeline

##### NMF model

To extract latent epigenetic patterns, we employed non-negative matrix factorization (NMF). Given a non-negative feature matrix $X \in \mathbb{R}_{\geq 0}^{n \times d}$, where $n$ is the number of samples (128263 CDSs) and $d$ is the number of features (20 epigenetic features), NMF factorizes

$X$ into two non-negative matrices: a weight matrix $W \in \mathbb{R}_{\geq 0}^{n\times r}$ and a pattern matrix $H \in \mathbb{R}_{\geq 0}^{d\times r}$, where $r$ is the dimensionality of the hidden space (Supplementary Fig. 1a). In our biological context, each column $\boldsymbol{h}_j$ of $H$ represents an epigenetic pattern defined by co-localization of epigenetic features, while each row $\boldsymbol{w}_i$ of $W$ quantifies the degree to which each pattern contributes to CDS $i$. This factorization is formulated as the following optimization problem:

$$\min_{W,H} \|X - WH^T\|_F^2,$$
$$\text{subject to } W \geq 0,\ H \geq 0,\ \|\boldsymbol{h}_j\|_1 = 1,$$

where $\boldsymbol{h}_j$ denotes the $j$-th column of $H$, and $\|\cdot\|_F$ is the Frobenius norm. The sum-to-one constraint on the columns of $H$, $\|\boldsymbol{h}_j\|_1 = 1$, is applied to resolve the scaling ambiguity inherent to NMF. This ambiguity arises because, for any arbitrary full-rank diagonal matrix $A$, the transformed pair, $\bar{W} = WA$ and $\bar{H} = HA^{-1}$, yields the same product $WH^{\mathrm{T}}$. To ensure comparability across features, we applied min-max normalization to the input matrix prior to decomposition.

#### Uniqueness of NMF solution

While increasing $r$ allows extraction of more complex patterns, it also raises concerns regarding the uniqueness and reproducibility of the NMF solution. Uniqueness here implies that any optimal solution $(\hat{W},\ \hat{H})$ satisfies $\hat{W} = W\Pi$ and $\hat{H} = H\Pi$, where $\Pi$ is a permutation matrix. According to theoretical work, uniqueness of $W$ and $H$ is guaranteed when the factor matrices are sufficiently scattered. Because verifying the sufficiently scattered condition is NP-hard, we implemented three empirical strategies to access solution reliability and to determine an appropriate value for $r$.

##### *Sparsity-based assessment*

First, we evaluated sparsity across replicates, based on the observation that if each column of $W$ and $H$ contains at least $r-1$ zero entries, the sufficiently scattered condition holds with high probability [51]. To avoid arbitrary thresholding, we extracted the $(r-1)$-th smallest value from each column and used the maximum among these values as an indicator. This process was repeated across 50 different initializations. We found that although the maximum $(r-1)$-th smallest values in $W$ remained close to zero for all values of $r$, the corresponding values in $H$ increased as $r$ increased, suggesting a loss of sparsity in the pattern matrix at higher dimensions (Supplementary Fig. 1b).

##### *Pattern Consistency Across Initializations*

Second, we tested whether consistent epigenetic patterns emerged across different initializations. We selected one solution $H^0 = [\boldsymbol{h}_1^0, \ldots \boldsymbol{h}_r^0]$ as a baseline and compared it with other 49 replicates $H^m = [\boldsymbol{h}_1^m, \ldots \boldsymbol{h}_r^m]$ $(m = 1, \ldots, 49)$. For each pair $(H^0, H^m)$, we iteratively matched columns based on maximum cosine similarity without duplication. The number of matched pairs exceeding predefined similarity thresholds (0.7, 0.8, or 0.9) was used to quantify reproducibility. When $r > 5$, the number of consistently matched patterns fell below $r$, indicating that not all patterns were robustly obtained. However, the number of consistent matches increased with $r$ even under the strictest threshold (0.9), which suggests that reproducible, non-random patterns continued to emerge (Supplementary Fig. 1c).

##### ***Weight matrix stability***

Finally, we assessed the stability of the weight matrix $W$ under different optimization strategies. Specifically, we compared the $W$ matrix obtained via joint optimization of $W$ and $H$ to that obtained by optimizing only $W$ with a pre-optimized $H$. The mean absolute error (MAE) between these two $W$ matrices increased sharply when $r > 15$, indicating instability in the weight assignment beyond this point (Supplementary Fig. 1d).

##### Selection of latent dimension and biological interpretability

Based on the sudden decline in reproducibility and stability beyond $r = 15$, we selected $r = 15$ as the optimal number of latent patterns. According to the sparsity-based analysis (Supplementary Fig. 1b), elements below $7.20 \times 10^{-2}$ were considered zero to satisfy the sufficient scattering condition [51]. Using this threshold (Supplementary Fig. 1g), the extracted patterns contained an average of $2.33 \pm 1.40$ features per pattern (mean $\pm$ SD; Supplementary Fig. 1h). This result suggests that NMF effectively capture feature combinations. Conversely, each feature contributed to $1.75 \pm 0.99$ patterns on average (Supplementary Fig. 1i), indicating that epigenetic features were shared across multiple patterns. Notably, repressive histone marks (H3K27me1, H3K27me3, H3K9me1, and H3K9me2) were predominantly assigned to distinct patterns (patterns 7 and 10; Supplementary Fig. 1j). This observation supports their context-dependent co-localization, consistent with prior findings [24]. The correlation distribution of patterns was skewed toward zero compared to that of epigenetic features (Supplementary Fig. 1e, f).

### Clustering analysis

#### CDS clustering

We classified CDSs based on the weight matrix $W$, which represents the degree to which each epigenetic pattern contributes to individual CDSs. Clustering was performed using the $k$-means algorithm, and the number of clusters was optimized to six using the elbow method implemented in the `yellowbrick` package (Supplementary Fig. 2a). These six clusters exhibited distinct average epigenetic pattern weights (Fig. 1d) and epigenetic feature profiles (Supplementary Fig. 2b), as well as diverse distributions of individual epigenetic pattern weights (Supplementary Fig. 2c). An UpSet plot further visualized the distribution of clusters across genes (Supplementary Fig. 2d).

##### Robustness of the clustering result

To evaluate the robustness of the clustering outcome, we systematically assessed how the removal of epigenetic patterns affected clustering accuracy. At each step, the epigenetic pattern whose removal resulted in the smallest decrease in accuracy was sequentially eliminated, using the original six-cluster result as the ground truth. Clustering accuracy remained above 80% as long as at least six specific patterns — namely patterns 13, 3, 8, 6, 9, and 12 — were retained (Supplementary Fig. 3a). We further assessed the impact of reducing the latent dimensionality by decreasing $r$ from 15 to 1. For each reduced $r$, we matched the resulting $r$ patterns to the original 15 patterns using cosine similarity, assigning zeros to unmatched components (i.e., the $15 - r$ unmatched patterns). We observed no clear relationship between clustering-relevant patterns and their reproducibility or appearance (averaged similarity) at smaller $r$ (Supplementary Fig. 3b). Consistently, clustering based on the six selected patterns (13, 3, 8, 6, 9, and 12) achieved over 80% accuracy, whereas using $r = 6$ directly yielded clustering results with lower accuracy (Supplementary Fig. 3c).

#### Statistical test for mutation rate differences between clusters

We tested whether the clusters exhibited statistically distinct mutation rates. Assuming mutations occur independently at each base with probability $\theta$, the number of mutations $x_k$ in a CDS $k$ of length $n_k$ follows a binomial distribution:

$$x_k \sim \mathrm{Bin}(n_k, \theta).$$

Let $\Lambda_i$ denote the set of CDSs in cluster $i$, and let $\theta_i$ be the mutation probability for CDSs in cluster $i$. We tested the following hypothesis:

- Null hypothesis (H0): $\theta_i = \theta_j$; all $x_k$ follow $\mathrm{Bin}(n_k, \theta_i)$ for $k \in \Lambda_i \cup \Lambda_j$.
- Alternative hypothesis (H1): $\theta_i \neq \theta_j$; $x_k \sim \mathrm{Bin}(n_k, \theta_i)$ for $k \in \Lambda_i$, and $x_k \sim \mathrm{Bin}(n_k, \theta_j)$ for $k \in \Lambda_j$.

The log-likelihoods for the two methods ($\ln(\mathcal{L}_0)$ and $\ln(\mathcal{L}_1)$) were given by:

$$\begin{aligned}
\ln(\mathcal{L}_0) &= \sum_{k \in \Lambda_i \cup \Lambda_j} \ln(\mathcal{L}(\theta_i \,|\, n_k, x_k)) = \sum_{k \in \Lambda_i \cup \Lambda_j} \ln \binom{n_k}{x_k} + x_k \ln(\theta_i) + (n_k - x_k) \ln(1 - \theta_i), \\
\ln(\mathcal{L}_1) &= \sum_{k \in \Lambda_i} \ln(\mathcal{L}(\theta_i \,|\, n_k, x_k)) + \sum_{k \in \Lambda_j} \ln(\mathcal{L}(\theta_j \,|\, n_k, x_k)) \\
&= \sum_{k \in \Lambda_i} \ln \binom{n_k}{x_k} + x_k \ln(\theta_i) + (n_k - x_k) \ln(1 - \theta_i) \\
&\quad + \sum_{k \in \Lambda_j} \ln \binom{n_k}{x_k} + x_k \ln(\theta_j) + (n_k - x_k) \ln(1 - \theta_j).
\end{aligned}$$

We computed the $p$-values using a log-likelihood ratio test with one degree of freedom, applying Bonferroni correction for multiple comparisons (Fig. 1e, g). In addition, we calculated McFadden's pseudo-$R^2$ as $1 - \frac{\ln(\mathcal{L}_1}{\ln(\mathcal{L}_0)}$ (Fig. 1h).

#### Clustering without NMF

To determine whether the observed mutation rate differences were attributable to latent epigenetic patterns or merely to the original epigenetic features, we also applied $k$-means clustering directly to the min-max normalized feature matrix $X$, bypassing NMF. The elbow method suggested five clusters, but we set the number to six to enable direct comparison with the NMF-based clustering (Fig. 1f). Notably, cluster 1 from the NMF-based results was split into clusters A and B, while clusters 2 and 6 were merged into cluster C, even though the latter two had distinct mutation rates. As a result, although clusters A and B showed significantly different mutation probabilities, the difference between clusters B and C was not statistically significant. This led to a total of five mutation rate-distinct clusters under the feature-based approach (Fig. 1g). These findings are consistent with the reduced pseudo-$R^2$ observed when clustering without NMF (Fig. 1h).

##### Chromatin state comparison

Chromatin states defined by ChromHMM [17, 19] are typically assigned to fixed-length genomic windows — e.g., 150 bp 150 bp [20] or 400 bp [21] — rather than to gene-based units such as CDSs. To enable comparison with our CDS-based clustering, we adapted these annotations by assigning chromatin states to each CDS. When a CDS spanned multiple states, it was segmented at state boundaries, and comparisons were made based on these modified units. Our analysis employed chromatin state annotations defined in two recent studies [20, 21], using the functional interpretations provided in [49]. The ChromHMM-based groupings showed minimal overlap with our clusters (Supplementary Fig. 4).

#### Explanatory power evaluation

##### Effects of classical genomic categories

We evaluated whether classical genomic categories could explain the variability in mutation probabilities observed across epigenetic clusters. CDSs were stratified based on the following features: CDS length, relative distance to the transcription start site (TSS), genomic position along chromosomes, expression level, essentiality, and lethality. For the Boolean categories (essentiality and lethality), CDSs were divided into two groups. For continuous variables, CDSs were divided into four quartile-based groups according to the value distribution of each feature. For each stratification, pseudo-$R^2$ scores were calculated under the assumption that mutation probabilities differed across clusters within each group. As a baseline, pseudo-$R^2$ scores were also computed using chromosome number as the grouping factor (Fig. 2a). Furthermore, we constructed two types of models: in one, mutation probabilities were defined solely by the group; in the other, they were defined by the combination of group and cluster. This resulted in 12 model variants for Boolean categories (2 groups $\times$ 6 clusters) and 24 variants for continuous categories (4 groups $\times$ 6 clusters) (Fig. 2b; Supplementary Fig. 5a). Although classical genomic features were distributed differently across clusters — such as in their group-wise distributions (Fig. 2c; Supplementary Fig. 5b, c) or the proportion of CDSs associated with essential (Supplementary Fig. 5d) and lethal (Supplementary Fig. 5e) genes — the statistical differences in mutation probabilities across clusters persisted (Fig. 2d; Supplementary Fig. 5f).

##### Signature of selection

To assess potential signatures of natural selection, we categorized SNVs using the Variant Effect Predictor (VEP) [52]. Gene annotations from the TAIR10 reference genome were used for transcript models (parameter `--gff`) and the genome sequence (parameter `--fasta`). All analyses were conducted using the Docker image `ensemblorg/ensembl-vep` available on Docker Hub. Among the possible VEP-defined categories of SNVs in coding regions — such as start retained, start lost, synonymous, missense, stop gained, splice donor, splice acceptor, stop lost, stop retained, coding sequence variant, and incomplete terminal codon — seven were observed in our data. To simplify the analysis, splice region variants, which were often co-labeled with other categories, were omitted. As a result, six categories were retained for downstream analysis: start lost, synonymous, missense, stop gained, stop lost, and stop retained (Supplementary Fig. 5g). Importantly, even when only synonymous variants were considered — thereby minimizing confounding from functional constraint — differences in mutation probability across clusters were still observed (Supplementary Fig. 5h).

##### Gene ontology analyses

Gene ontology (GO) enrichment analysis was performed using the `clusterProfiler` package (v4.4.4) in R, with the `org.At.tair.db` database (v3.15.1) and the biological process ontology. A Bonferroni-adjusted $p$-value threshold of 0.01 was applied to identify significantly enriched GO terms. For the analysis, only genes whose CDSs were exclusively assigned to a single cluster were considered. To improve interpretability, redundant GO terms were simplified using the default simplification procedure (Supplementary Fig. 6).

#### Mutation rate estimation for individual CDSs

##### GLM model for epigenetic pattern weights

To estimate the mutation probability for each CDS, we applied a generalized linear model (GLM) under the assumption of binomially distributed mutation counts. Specifically, we posited that the mutation probability $\theta_k$ of a CDS $k$ is determined by its epigenetic-pattern weights $\boldsymbol{w}_k$, corresponding to the $k$-th row of the weight matrix $W$. The GLM was formulated with the logit function as the link function, yielding the following model:

$$
\begin{aligned}
x_k &\sim \mathrm{Bin}(n_k, \theta_k), \\
\mathrm{Logit}(\theta_k) &= \ln\left(\frac{\theta_k}{1-\theta_k}\right) = \boldsymbol{\beta}^{\mathrm{T}}\boldsymbol{w}_k + \beta_0, \\
\theta_k &= \frac{1}{1+\exp(-\boldsymbol{\beta}^{\mathrm{T}}\boldsymbol{w}_k + \beta_0)},
\end{aligned}
$$

where $x_k$ is the number of observed mutations in CDS $k$, $n_k$ is its length in base pairs, $\boldsymbol{\beta}$ is the vector of regression coefficients associated with the epigenetic patterns, and $\beta_0$ is the intercept term. Both $\boldsymbol{\beta}$ and $\beta_0$ were estimated via maximum likelihood. The z-values and corresponding Wald test $p$ values for individual epigenetic patterns were also obtained from the fitted GML (Fig. 2f). Also, the distribution of predicted probabilities were compared across clusters with a one-sided Mann-Whitney $U$ test for all pairs of clusters $i$ and $j$ ($i < j$) to evaluate the observed decay in clusters are reserved (Fig. 2g).

##### Statistical power evaluation

To assess the explanatory power of the epigenetic-pattern-based GLM, we computed the log-likelihood of the fitted model (alternative model) based on the predicted mutation probabilities $\theta_k$. This was compared against a null model in which all CDSs were assumed to share a common mutation probability. We further extended the GLM framework to compare the predictive power of other types of features — such as classical gene categories and 3-mer sequence content — by constructing alternative models in which $\theta_k$ was modeled as a function of those features. These models were evaluated against their respective null models using McFadden's pseudo-$R^2$ as a measure of explanatory power. Pseudo-$R^2$ values and $z$-values across models with different feature sets were compared to assess their relative contributions to mutation rate variation (Fig. 2e; Supplementary Fig. 7).

### Mutation accumulation lines

We obtained a published SNV dataset from A. thaliana mutation accumulation lines generated via single-seed descent [15]. To focus on likely de novo mutations, we excluded SNVs that were shared across multiple lines. Although the NMF-derived clusters exhibited variation in mutation probabilities, no statistically significant differences were detected by the log-likelihood ratio test (Supplementary Fig. 8a). Nevertheless, a consistent trend was observed: CDSs assigned to clusters 5 and 6 showed lower mutation probabilities than those in clusters 1 and 2 (Fig. 3a), indicating that the cluster-level mutation trends persist even in germline-derived mutation datasets.

### Hypoxia analysis

#### Datasets and processing

Raw FASTQ files for H3K14ac and H3K9ac under 9-hour hypoxia stress and corresponding control conditions were obtained from GSE122804. All preprocessing steps followed the original pipeline [27]. Barcode sequences were trimmed using `fastx_clipper`, and short or low-quality reads were filtered using `fastq_quality_filter` with parameters `-q 20` and `-p 80`. Filtered reads were aligned to the TAIR10 reference genome using Bowtie2. Bigwig files were generated using `bamCoverage` with the `-normalizeUsing PRKM` option and the black list provided in the original study. For gene expression, we additionally used the fold-change values of transcript levels under 9-hour hypoxia stress from the same study [27]. For our analysis, we averaged replicate BigWig signals and assigned the resulting values to individual CDSs. Hypoxia-responsive genes (HRGs) were defined as the 49 core HRGs established in previous studies [27, 28], supplemented by five evolutionarily conserved group VII ethylene response factor (ERFVII) genes: RAP2.2, RAP2.3, RAP2.12, HRE1, and HRE2. Since HRE2 appears in both lists, the final set comprised 53 unique hypoxia-responsive genes.

#### Pattern-14 activation scores

To capture the joint activation of H3K9ac and H3K14ac — histone marks comprising epigenetic pattern 14 (Fig. 1b), which is associated with increased mutation probability (Fig. 2f) — we computed the fold changes of each acetylation mark at the CDS level. The lower of the two values was taken as the pattern-14 activation score for each CDS. Under the original dataset utilized to generate epigenetic pattern matrix, the lowest values of H3K9ac, H3K14ac, and pattern-14 weights were positively correlated in higher value ranges (Supplementary Fig. 8b), supporting the validity of using this simple score as a proxy for pattern-14 activation.

#### Pathway analysis

To interpret the functional significance of pattern-14 activation at the gene level, we performed Gene Set Enrichment Analysis (GSEA) using the `gseGO` function from the `clusterProfiler` R package with the biological process ontology. Genes were ranked by their maximum pattern-14 activation score across all CDSs, and enrichment was assessed using this ranked gene list (Fig. 3c). For comparison, GSEA was also performed using genes ranked by expression fold changes under hypoxia (Supplementary Fig. 8d). Pattern-14-based ranking yielded stronger enrichment for the DNA damage response

(GO:0006974) than the expression-based ranking (NES = 1.78 vs 1.56; adjusted $p = 6.51 \times 10^{-8}$ vs $3.33 \times 10^{-4}$; Fig. 3c). The similar trend was observed for its key subclass, DNA repair (GO:0006281), which also showed higher enrichment in the pattern-14-based GSEA (NES = 1.77 vs 1.55; adjusted $p = 5.16 \times 10^{-7}$ vs $7.81 \times 10^{-4}$). Motivated by this difference, we further analyzed CDS-level activation scores using gene-to-GO annotations from the TAIR10 database. For each GO term, we tested whether the distribution of pattern-14 activation scores among associated CDSs differed significantly from the background, using a two-sided Mann-Whitney $U$ test, followed by Bonferroni correction. This CDS-level enrichment analysis revealed significant increases in pattern-14 activation among CDSs assigned with DNA repair (GO:0006281) and its subpathways, including double-strand break repair (GO:0006302), nucleotide-excision repair (GO:0006289), recombinational repair (GO:0000725), and single strand break repair (GO:0000012) (Fig. 3d). These enrichments were not detected when analyzing expression fold changes (Supplementary Fig. 8e).

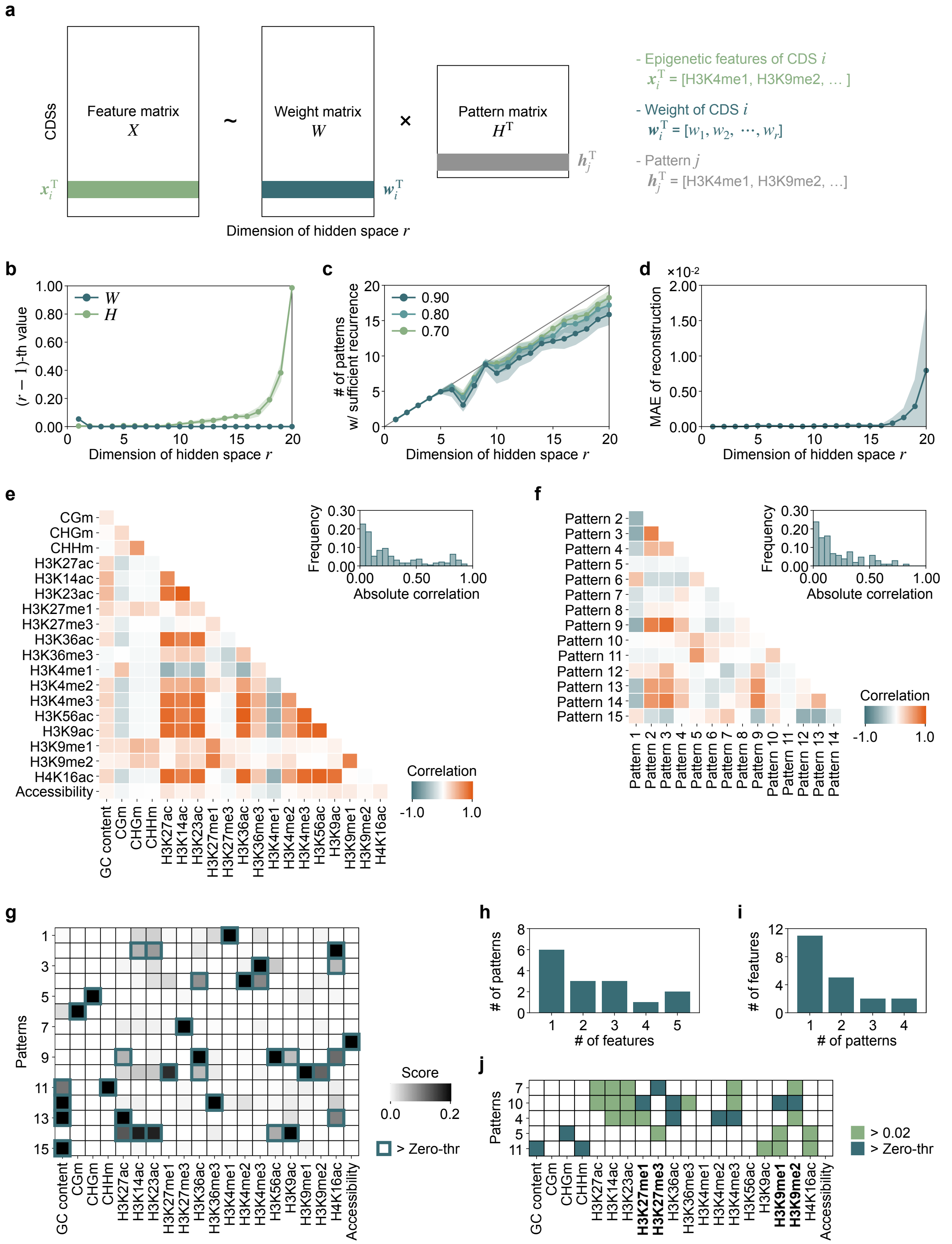


**Supplementary Fig. 1. Implementation of NMF.**

**(a)** Setup of NMF, where the epigenetic feature matrix $X$ is decomposed into two matrices: the weight matrix $W$ and the pattern matrix $H$. **(b)** Largest value of the $(r-1)$ -th smallest element in each column of $W$ or $H$ (mean ± SD over 50 trials). **(c)** Number of patterns surpassing the cosine similarity criterion (0.90, 0.80, or 0.70; mean ± SD over 49 trials). **(d)** MAE between $W$ optimized simultaneously with $H$ and $W$ optimized using the pre-optimized $H$ (mean ± SD over 50 trials). **(e, f)** Pearson correlation coefficients across epigenetic features (a) and epigenetic pattern weights (b). Insets show absolute coefficients. **(g)** Extracted epigenetic pattern matrix. Dark green outlines enclose areas with values greater than the zero-threshold determined by (b). **(h)** Histogram showing the distribution of the number of features within each pattern. **(i)** Histogram showing the distribution of the number of patterns in which each feature appears. **(j)** Patterns consisting of repressive markers. Dark green areas indicate values larger than the zero-threshold, light green areas indicate values larger than 0.02, and bolded features represent repressive markers.

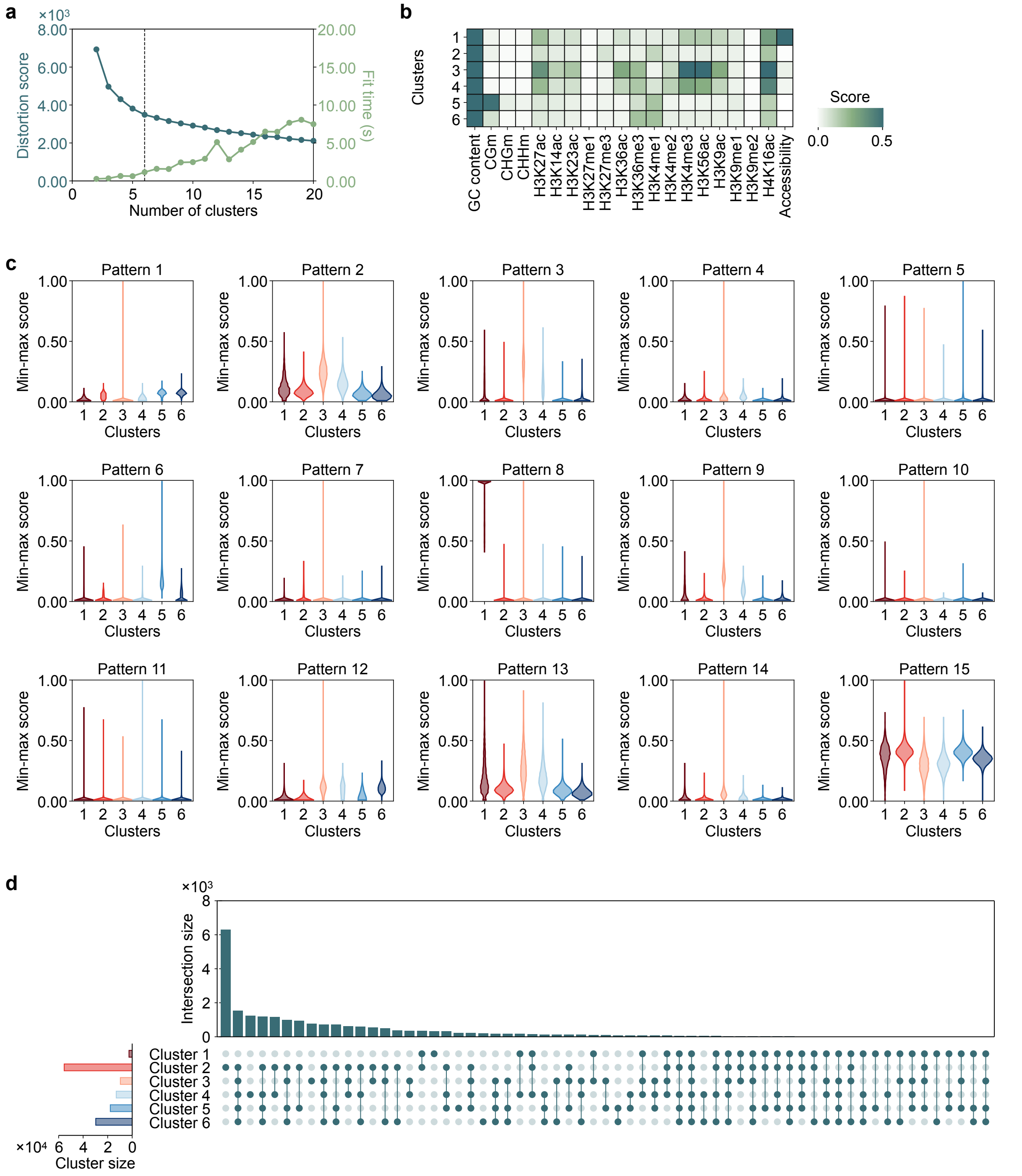


**Supplementary Fig. 2. Clustering results.**

**(a)** Results of the elbow method. The black line marks the optimal number of clusters. **(b)** Heatmap showing the average weights of epigenetic features across clusters. **(c)** Distribution of epigenetic pattern weights. **(d)** UpSet plot showing the composition of clusters among genes.

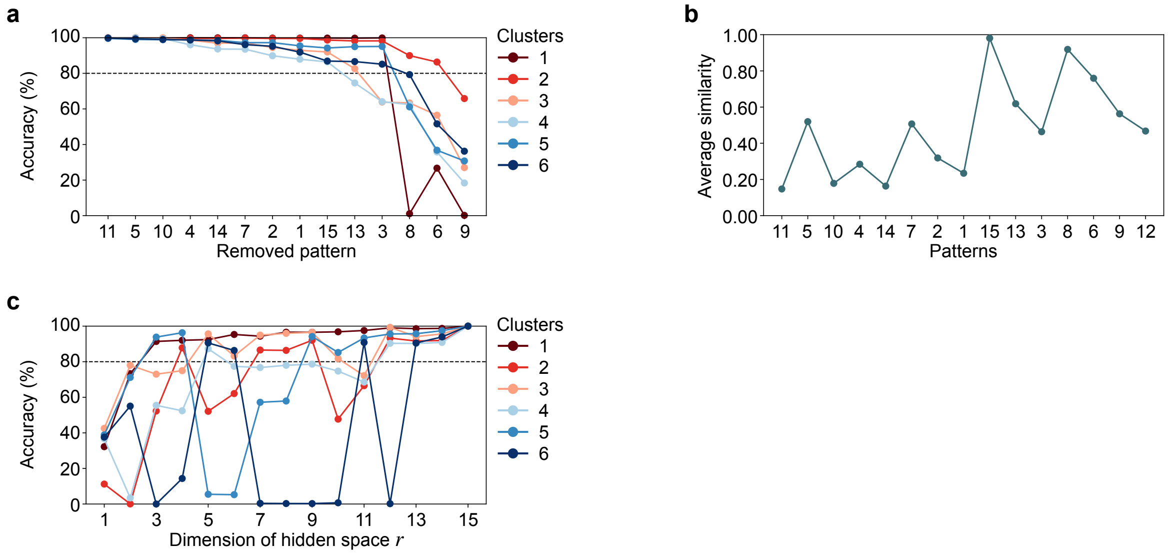


**Supplementary Fig. 3. Effect of epigenetic patterns on clustering results.**

**(a)** Impact of the sequential removal of patterns on clustering accuracy. **(b)** Average cosine similarity of patterns across hidden space dimensions ($r = 1, \ldots, 15$). **(c)** Clustering accuracy based on NMF results for different hidden space dimensions r.

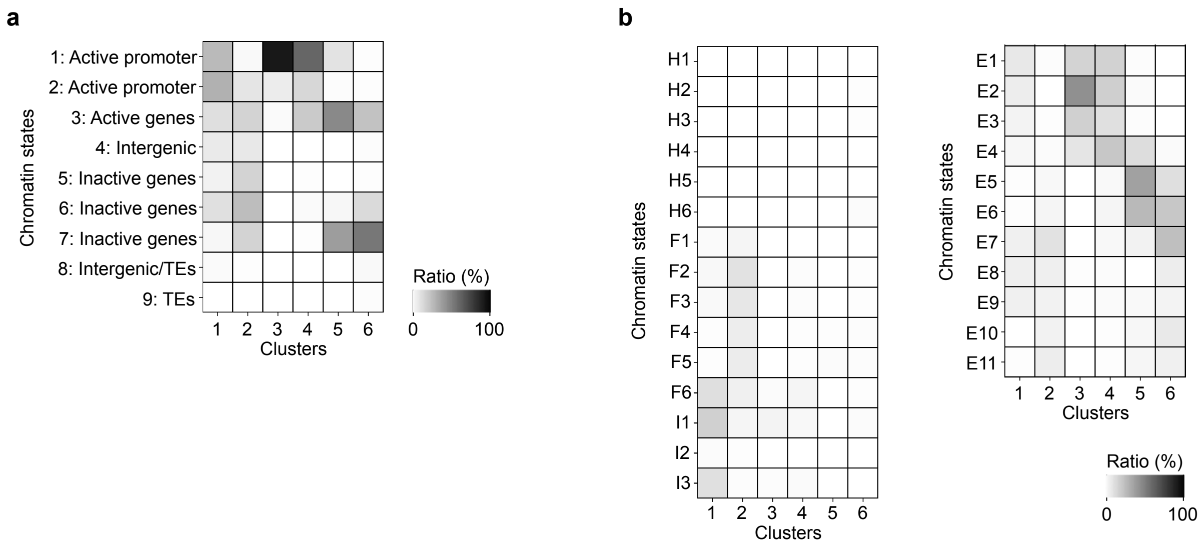


**Supplementary Fig. 4. Comparison to the chromatin states.**

**(a, b)** Results of comparison with chromatin states from Sequeira-Mendes et al. [20], and annotations by Quadrana et al. [49] (a), and from Jamge et al. [21] (b).

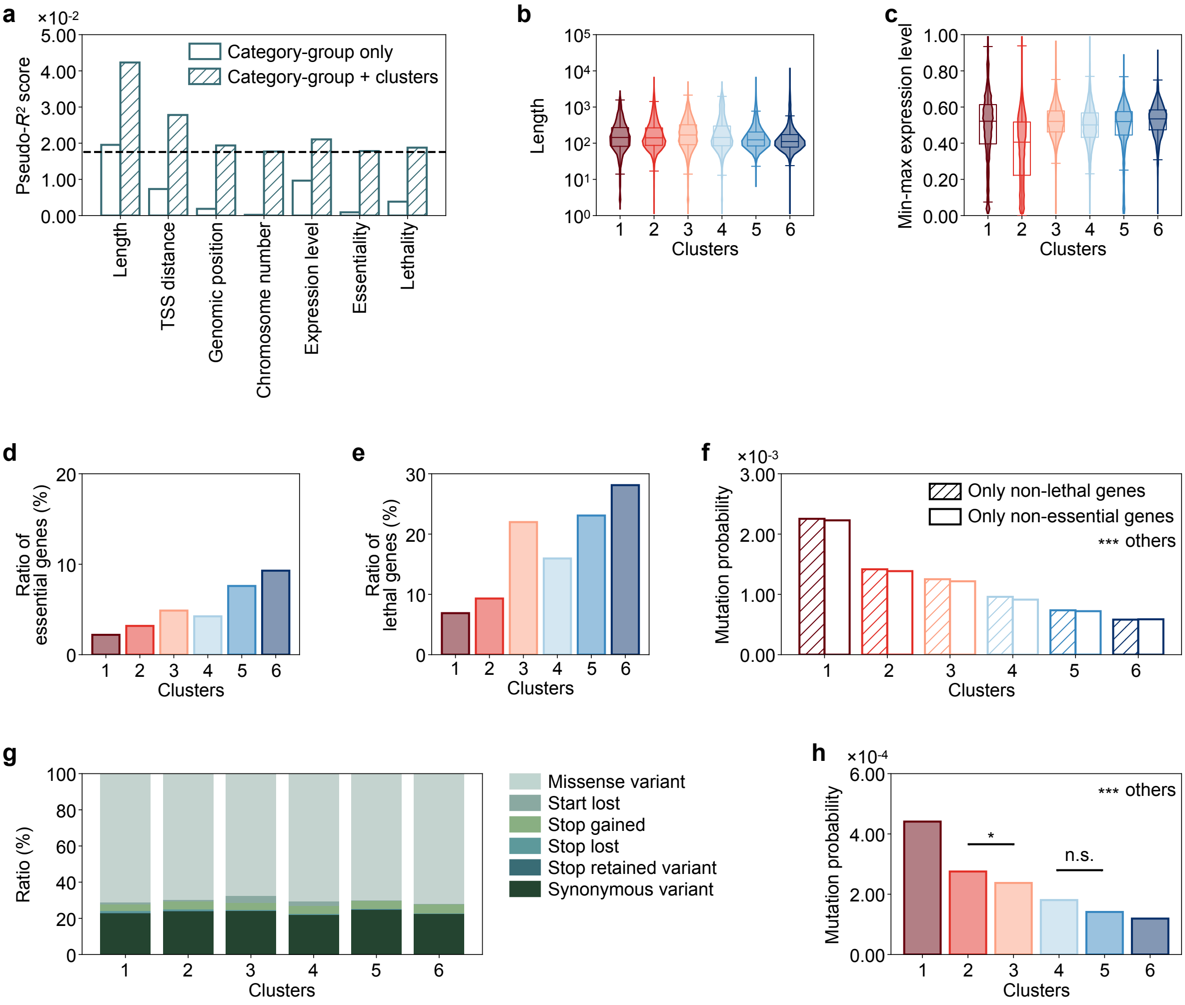


**Supplementary Fig. 5. Effects of classical genomic categories.**

**(a)** Pseudo-$R^2$ score for mutation probabilities explained by groups defined by each classical genomic category and by the combination of groups and clusters. **(b)** Distribution of CDS lengths across clusters. **(c)** Distribution of min-max normalized expression levels across clusters. **(d, e)** Ratio of CDSs included in lethal genes (d) and essential genes (e). **(f)** Mutation probabilities of clusters calculated using only CDSs from non-lethal or non-essential genes. For all cluster pairs generated using only non-lethal genes and for all cluster pairs generated from non-essential genes, the log-likelihood ratio test yielded $***p < 0.001$ after Bonferroni correction. **(g)** Composition of variant types across clusters. **(h)** Mutation probabilities of clusters calculated using only synonymous variants. For all cluster pairs, unless otherwise noted ($*p < 0.05$; n.s., not significant), the log-likelihood ratio test yielded $***p < 0.001$ after Bonferroni correction.

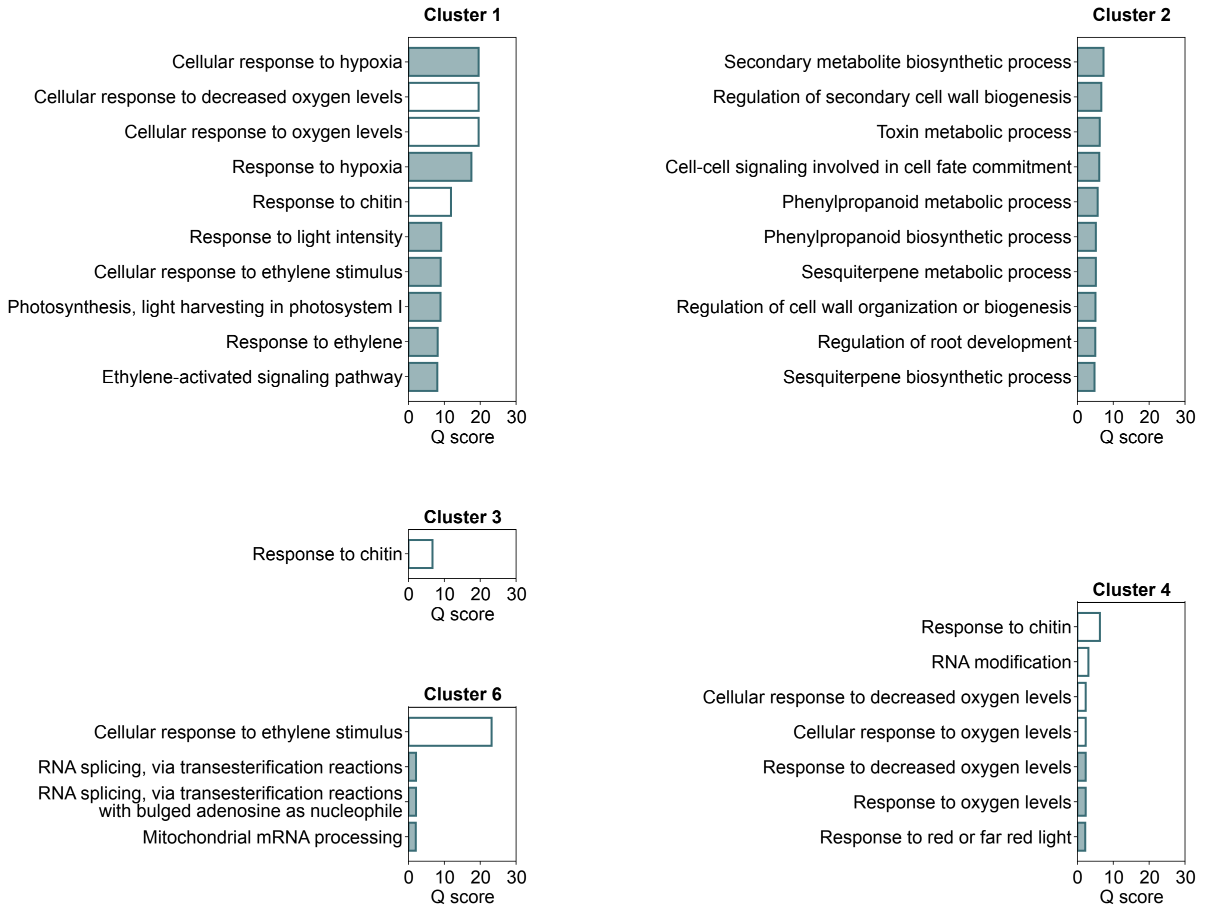


**Supplementary Fig. 6. GO terms for genes composed exclusively of CDSs from a specific cluster.**

The white bars represent GO terms shared with other clusters, while the green bars represent GO terms unique to each cluster.

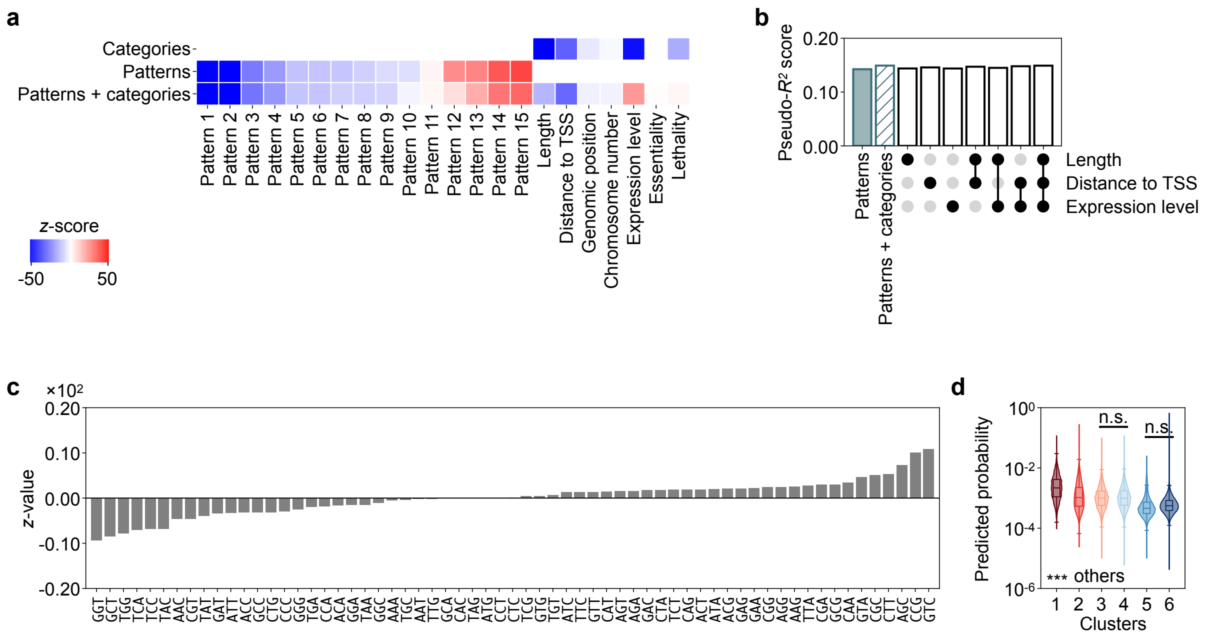


**Supplementary Fig. 7. Explanatory power of various features.**

**(a)** Heatmap showing $z$-scores in the GLM prediction using classical genomic categories, patterns, and both. **(b)** Pseudo-$R^2$ score for mutation probabilities explained by various combinations. The included features are indicated by filled circles in the matrix below the bars. **(c)** $z$-scores in the GLM prediction using 3-mer context. **(d)** Distribution of predicted probabilities among clusters using categories and 3-mer in addition to patterns. For all pairs of Clusters $i$ and $j$ ($i < j$), unless otherwise noted (n.s., not significant), a one-sided Mann-Whitney $U$ test yielded $^{***}p < 0.001$ after Bonferroni correction.

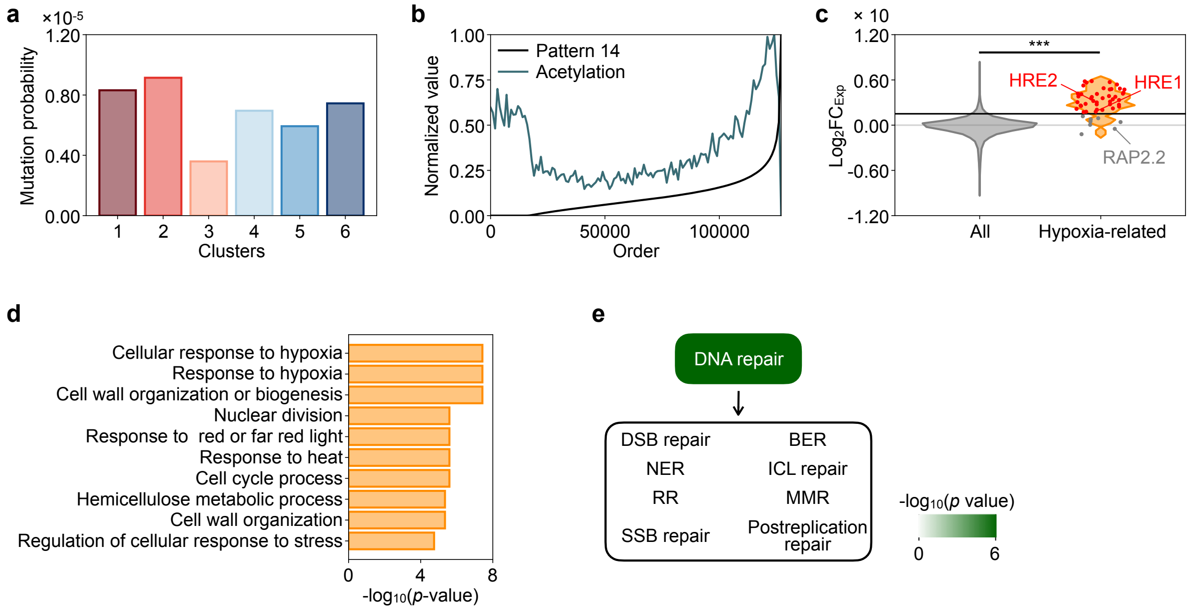


**Supplementary Fig. 8. Expression up-regulation at fitness-relevant loci.** **(a)** MA line-based mutation probabilities of CDSs. No cluster pairs showed statistical significance in the log-likelihood ratio test after Bonferroni correction ($p > 0.05$). **(b)** Normalized pattern-14 weights and the maximum values of H3K9ac and H3K14ac in the original epigenetic datasets, sorted by pattern-14 weights. **(c)** Distribution of expression up-regulation for all genes and for hypoxia-related genes, with data points in the top 5% highlighted in red. Selected points are labeled with gene names. A one-sided Mann-Whiney $U$ test revealed $^{***}p < 0.001$. **(d)** Top 10 GO terms identified by GSEA, ranked by expression up-regulation. **(e)** Enrichment analysis based on the Mann-Whitney $U$ test $p$-values after Bonferroni correction for genes belonging to each GO term compared with the background distribution of expression up-regulation. Green shading indicates $p$-values, while GO terms in the white area show no significant enrichment ($p > 0.05$).
